# Membrane curvature sensing of the lipid-anchored K-Ras small GTPases

**DOI:** 10.1101/492710

**Authors:** Hong Liang, Huanwen Mu, Frantz Jean-Francois, Bindu Lakshman, Suparna Sarkar-Banerjee, Yinyin Zhuang, Yongpeng Zeng, Weibo Gao, Dwight V. Nissley, Alemayehu A. Gorfe, Wenting Zhao, Yong Zhou

## Abstract

Cell morphologies, defined by plasma membrane (PM) local curvature, change during mitogen-dependent function and pathology, such as growth, division and proliferation. The lipid-anchored Ras small GTPases are essential upstream regulators of the mitogen-activated protein kinases (MAPKs) cascades and play key roles in many pathological conditions, especially cancer. Ras signaling is mostly compartmentalized to the cell PM through the formation of nanometer-sized domains, termed as nanoclusters, and undergo selective lipid sorting for efficient effector recruitment and activation. Thus, Ras function might be sensitive to changing PM curvature, potentially regulating mechanosensing of mitogen signaling. We employed nanofabrication and super-resolution imaging and found that Ras functions respond to PM curvature modulations in an isoform specific manner: nanoclustering and signaling of the most oncogenically prevalent isoform K-Ras favor less curved PM, while those of another isoform H-Ras favor more curved PM. We then examined whether Ras membrane curvature sensing is mediated by lipid sorting. We found that anionic phospholipids sense changing PM curvature in distinct manners: phosphatidylserine (PS) localization shows preference for less curved membrane but phosphoinositol 4,5-bisphosphate (PIP_2_) localization favors more curved PM. Depletion of endogenous PS abolishes K-Ras PM curvature sensing. Exogenous PS addback and synthetic bilayer binding assays further show that only mixed-chain PS species, but not other PS species, mediate K-Ras curvature sensing. Taken together, the Ras proteolipid nano-assemblies on the PM act as relay stations to convert mechanical stimulations to mitogenic signaling circuits, thus a novel mechanism for cancer cell mechanotransduction

## Introduction

Cell morphology changes during mitogen-regulated growth, division, proliferation and migration ^1^, and correlates with mitogen-dependent cancer cell transformation and epithelial-mesenchymal transition ^2^. How cell morphology corresponds with the intracellular mitogenic signaling is poorly understood. Cell shape is defined by the changes in the plasma membrane (PM) curvature, which may alter the spatial distribution of lipids ^3^. The lipid-anchored Ras small GTPases (H-Ras, N-Ras, and splice variants K-Ras4A and K-Ras4B) are essential regulators of mitogenic signaling and are highly mutated in cancer ^4,5^. Ras signaling is mostly compartmentalized to the PM ^4–6^, which causes Ras function to be potentially sensitive to the changing lipid distribution at the curved PM. Ras isoforms use their C-terminal membrane anchoring domains to interact with PM constituents ^5,6^. Because of their structurally distinct membrane-anchoring domains, Ras isoforms interact with lipids in the highly heterogeneous PM in an isoform-specific manner ^6^. The selective lipid sorting of Ras results in the formation of nanometer-sized domains, termed as nanoclusters, in an isoform-dependent manner ^6,7^. Ras/lipid nanoclusters are essential to the signaling of both wild-type and constitutively active oncogenic mutants of Ras. This is because most of the Ras downstream effectors contain specific lipid-binding domains and require binding to both active Ras and specific lipid types for efficient PM recruitment and activation ^6,8^. Thus, the structural integrity of the Ras proteolipid nano-assemblies and the Ras-dependent MAPK signaling may depend on membrane curvature. To test this hypothesis, we manipulated the curvature of the PM of intact/live cells, isolated native PM or synthetic bilayers via established and novel methods. Using quantitative imaging methodologies, we show that the PM localization, the nanoclustering and the signaling of K-Ras4B (the most oncogenically prevalent isoform and simply referred to as K-Ras below) favor low curvature whereas those of H-Ras prefer large curvature. We further explored potential molecular mechanism and found that the membrane curvature sensing of K-Ras is mediated via the selective sorting of distinct anionic phospholipid phosphatidylserine (PS) species. Thus, Ras/lipid nanoclusters on the PM act as structural relay stations to convert cell surface morphological modulations to intracellular mitogenic signaling.

## Results

### Plasma membrane localization of Ras senses plasma membrane curvature modulations in an isoform specific manner

We first tested how the lipid-driven Ras spatiotemporal organization on the PM may be sensitive to PM curvature. Membrane curvature is a highly dynamic property difficult to manipulate because PM curvature is linked with many other membrane properties, such as mechanics, cytoskeletal organization and extracellular matrix. We, thus, employed a series of novel and established methodologies to manipulate PM curvature, in an effort to better correlate the membrane curvature-dependent phenomena. Specifically, in intact/live cells, we used a novel nanofabrication technique to precisely induce PM curvature in live cells ^9^, expression of Bin-amphiphysin-Rvs domains (BAR) domains to induce PM curvature ^10,11^, or incubation of cells/isolated native vesicles in buffers with different osmolality to introduce osmotic imbalance across the PM to modulate PM curvature ^12,13^. In synthetic vesicles, we used vesicle sizes to define bilayer curvature ^14,15^.

The first method employed in our study was a novel nanofabrication technology, which manufactures an array of nanobars with a length of 2µm and a width of 250nm (yielding a curvature radius of 125nm at the curved ends) on SiO_2_ substrates (Fig.1A) ^9^. Thus, each nanobar contains two highly curved ends and a comparably flat center region ^9^. *Homo sapiens* bone osteosarcoma U-2OS cells were grown over the nanobars, with their basolateral PM tightly wrapping around the nanobars and generating PM curvature with a precise curvature radius ^9^. We used confocal imaging to quantify the fluorescence distribution of GFP-K-Ras^*G12V*^ (a constitutively active K-Ras oncogenic mutant), GFP-tH (minimal anchor of H-Ras) or mCherry-CAAX motif on nanobars (Fig.1B-G). We focused on these 3 protein/peptides because they have been established to sort distinct lipid compositions ^16–19^. Specifically, K-Ras^*G12V*^ selectively sorts PS, while tH heavily favors cholesterol and phosphoinostol 4,5-bisphosphate (PIP_2_) ^16–18^. On the other hand, the CAAX motif interacts with the membranes randomly and lacks the ability to detect structural changes in the lipid bilayers ^9^. To better quantify the potential PM curvature preferences, we calculated the fluorescence ratio at curved ends / flat center as a measure of PM curvature preferences ^9^. Averaged fluorescence heatmaps and quantification show that mCherry-CAAX had the lowest fluorescence intensity ratio between the curved nanobar ends and the flat nanobar center (Fig.1H and I). GFP-K-Ras^*G12V*^ and GFP-tH distributed more to the curved ends, with GFP-tH displaying a higher curvature preference (Fig.1H and I). The frequency distribution further shows that the ends / center fluorescence ratios of GFP-tH distributed more towards the curved ends of the nanobars than those of GFP-K-Ras^*G12V*^ (Fig.1J). Taken together, the PM localization of Ras proteins senses membrane curvature changes in an isoform-specific manner: the PM binding of K-Ras^*G12V*^ favors less curved PM than H-Ras anchor, whereas the CAAX motif showed minimal preference for PM curvature.

**Figure 1.**
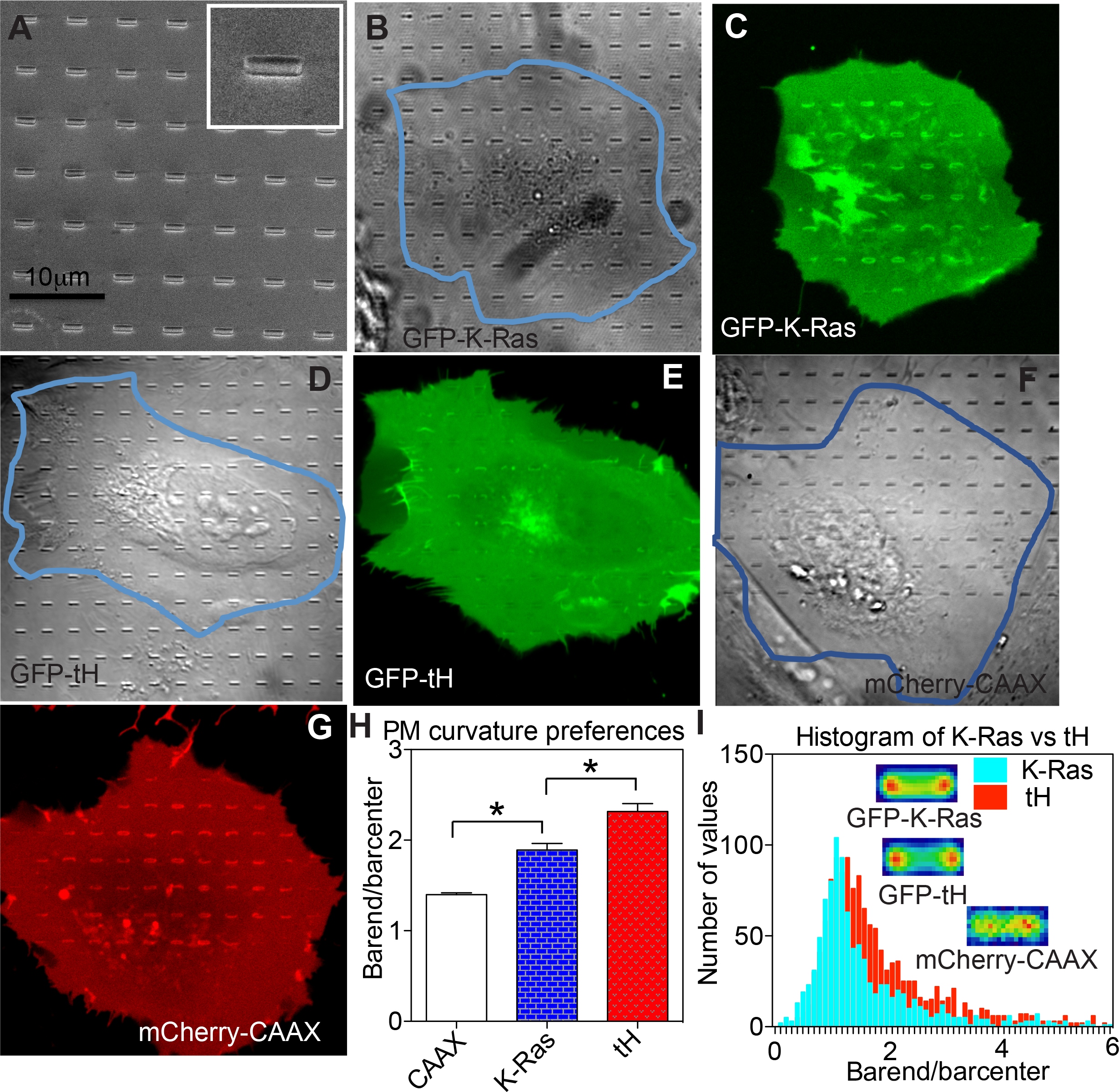
Ras localization to the cell PM senses curvature modulations in an isoform-specific manner. (**A**) A scanning EM image shows a SiO_2_ substrate etched with an array of nanobars with length of 2µm and width of 250nm. The inset shows a single nanobar. U-2OS cells expressing GFP-K-Ras^*G12V*^ grown over the nanobars are shown in phase contrast (**B**) and confocal (**C**), GFP-tH in phase contrast (**D**) and confocal (**E**), or mCherry-CAAX in phase contrast (**F**) and confocal (**G**). (**H**) Fluorescence intensity ratios between the curved ends and the flat center portions of the nanobars were calculated to indicate the preferential localization of various Ras protein/peptides on the basolateral PM. Data are shown as mean±SEM, with * indicating statistical significance p < 0.05 evaluated using the one-component ANOVA. (**I**) Frequency distribution of all the individual nanobar end/center fluorescence ratios of GFP-K-Ras^*G12V*^ or GFP-tH is shown. A total of 1007 nanobars for GFP-K-Ras^*G12V*^, 1377 nanobars for GFP-tH and 415 nanobars for mCherry-CAAX were imaged and caculated. The insets show compiled images of the averaged fluorescence intensities heatmaps of all the nanobars imaged.

To further validate the curvature-induced changes in Ras localization to the cell PM, we next performed electron microscopy (EM)-immunogold labeling ^17,18^. To better correlate with the nanobar data above, we used several established methods to modulate PM curvature: (1) to further induce PM curvature, we ectopically expressed the membrane curvature sensing/modulating BAR domain of amphiphysin 2 (BAR_amph2_) ^20,21^; (2) to decrease PM curvature, we incubated cells in hypotonic medium, where a typical growth medium was diluted with 40% deionized water, for 5 minutes. We, thus, established a gradient of PM curvature magnitudes: *<u>flat</u>* (hypotonic incubation), *<u>normal</u>* (untreated cells), *<u>elevated</u>* (expression of BAR domains). After acute PM curvature modulations, the apical PM sheets of baby hamster kidney (BHK) cells expressing GFP-tagged Ras were attached to copper EM grids and immunolabeled with 4.5nm gold nanoparticles conjugated to anti-GFP antibody. The PM sheets were imaged using transmission EM (TEM) at a magnification of 100,000x and the number of gold nanoparticles within a select 1µm^2^ PM area was counted as an estimate of PM binding of the GFP-tagged Ras (Fig.S1A, C and E). Fig.2A (as well as Fig.S1I) shows the mean number of gold particles/1µm^2^ PM area in BHK cells with the elevated PM curvature. The gold numbers of the full-length constitutively active and oncogenic mutant GFP-H-Ras^*G12V*^ (H-Ras) or that of the truncated minimal membrane-anchoring domain GFP-tH (tH) were markedly higher than those of the oncogenic mutant GFP-K-Ras^*G12V*^ (K-Ras) or the truncated minimal membrane-anchoring domain GFP-tK (tK) (Fig.2A and Fig.S1I). This data is entirely consistent with the nanobar data shown in Fig.1. To further validate the potential PM curvature preferences of Ras isoforms, we flattened BHK cell PM via incubating the cells in hypotonic medium for 5 minutes in the same batch of EM experiments. Fig.2B and Fig.S1I show that, with the PM flattened, the PM binding of GFP-K-Ras^*G12V*^ or GFP-tK is significantly higher than that of H-Ras constructs, also consistent with the observed Ras isoform-specific curvature preferences in Fig.1. Fig.2C and Fig.S1I show the control EM experiments, where the immunogold labeling was equivalent for all Ras constructs tested in untreated cells with normal extent of PM curvature.

**Figure 2.**
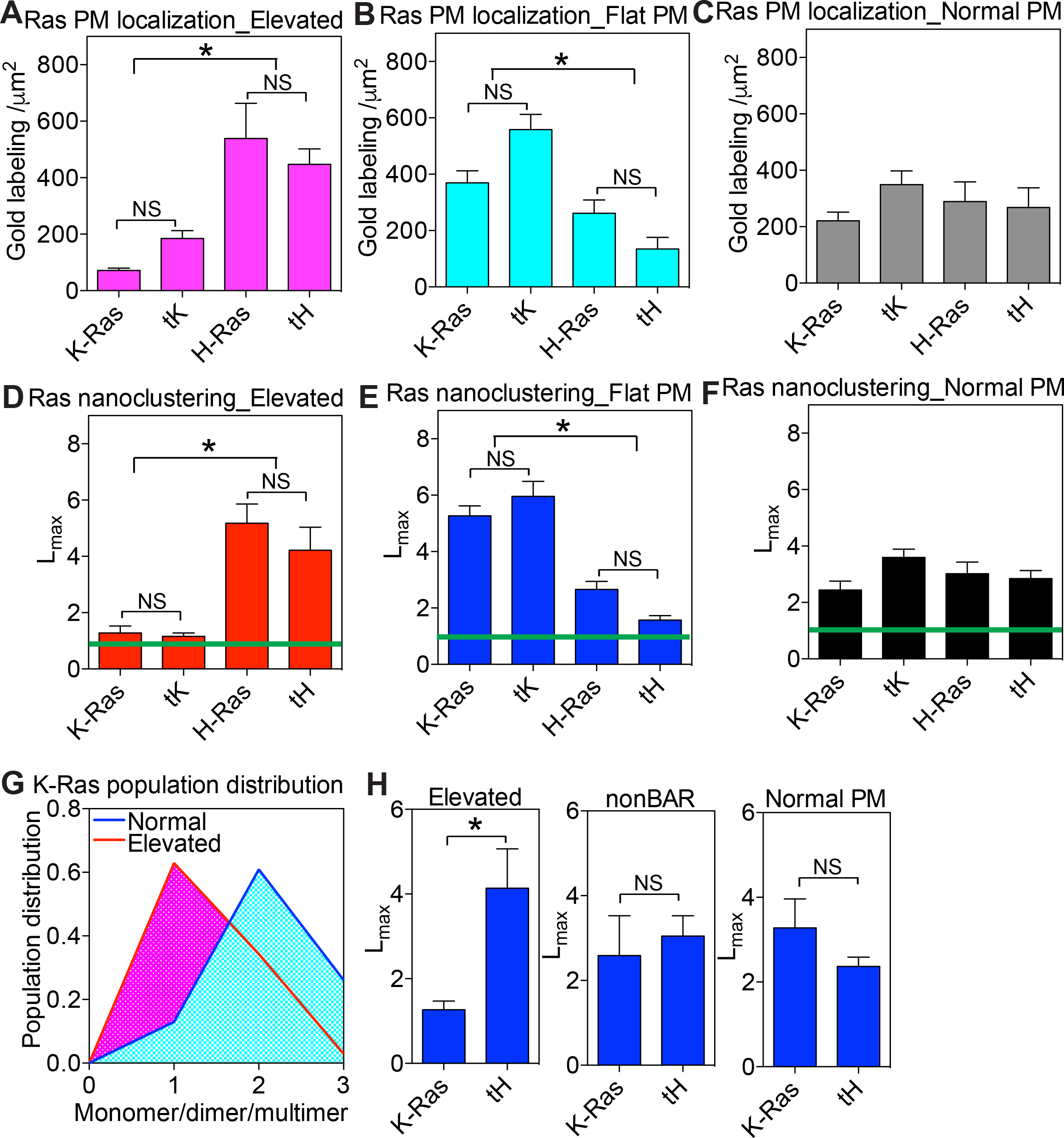
Ras spatiotemporal organization on the PM depends on PM curvature modulations. BHK cells were ectopically expressed with GFP-tagged Ras constructs (*K-Ras*: constitutively active oncogenic mutant K-Ras^*G12V*^; *tK*: the truncated minimal membrane-anchoring domain of K-Ras; *H-Ras*: the oncogenic mutant H-Ras^*G12V*^; *tH*: the truncated minimal membrane-anchoring domain of H-Ras). Intact apical PM sheets of the BHK cells were attached to EM grids and immunolabeled with 4.5nm gold nanoparticles conjugated to anti-GFP antibody. The number of gold particles per 1μm^2^ area on the intact PM sheets was counted to estimate the localization of the GFP-tagged Ras on the PM in cells with elevated PM curvature (**A**, local curvature induced by ectopic expression of BAR_amph2_ domain), flattened PM (**B**, local curvature decreased via a 5-minute incubation of cells in hypotonic medium further diluted with 40% deionized water), or with normal curvature (**C**, untreated cells). To analyze the lateral spatial distribution of the Ras proteins/peptides on the PM, the univariate K-function was used to quantify the nanoclustering of the gold-labeled GFP-Ras constructs in the same EM images used above. The optical nanoclustering, *L*_*max*_, for the GFP-Ras constructs in cells with elevated PM curvature (**D**), flattened PM (**E**), or with normal curvature (**F**). The green lines indicate the 99% confidence interval (99%C.I.). (**G**) The extent of lateral aggregation of GFP-K-Ras^*G12V*^ was evaluated using RICS analysis on the PM of BHK cells co-expressing GFP-K-Ras^*G12V*^ with either empty vector pC1 (normal curvature) or RFP-BAR_amph2_ (elevated curvature). (**H**) The nanoclustering (*L*_*max*_) of GFP-K-Ras^*G12V*^ or GFP-tH was calculated in BHK cells with elevated (ectopic expression of RFP-BAR_FCC_), nonBAR (ectopic expression of the truncated RFP-BAR_FCH_), or normal PM curvature (untreated cells). All data are shown as mean±SEM. In all the gold number counting calculations, one-component ANOVA was used to evaluate the statistical significance between various groups, with * indicating p < 0.05. In all the nanoclustering analyses, bootstrap tests compared the test samples with 1000 bootstrap samples and were used to evaluate the statistical significance between different sets, with * indicating p < 0.05.

### Ras nanoclustering/oligomerization depends on PM curvature modulations in an isoform specific manner

Once localized to the cell PM, Ras proteins selectively sort PM lipids to laterally form nanometer-sized domains, termed as nanoclusters, in an isoform-specific manner. Ras nanodomains concentrate select sets of lipids, which is functionally important because most Ras effectors contain specific lipid-binding domains and require binding to particular lipids for proper PM localization and activation. Ras nanoclustering does not necessarily correlate with their localization to the membranes. For instance, changing PS levels in the PM directly alters both the PM localization and the nanoclustering of K-Ras ^16,18,22,23^. On the other hand, altering transmembrane potential or actin organization perturbs only K-Ras lateral clustering without changing its PM localization ^18,24^. Supplementation of distinct PS species, such as the mono-unsaturated di18:1 PS (DOPS), increases K-Ras PM localization without changing its nanoclustering ^17^. To quantify the potential sensitivity of Ras nanoclustering to PM curvature modulations, we used the same electron micrographs utilized in the above gold number analyses (Fig.2A-C) and calculated the lateral spatial distribution of the immunogold labeling via the Ripley‘s K-function analysis ^17–19^. Fig.S1B, D and F display the color-coded spatial distribution of the gold nanoparticles in the corresponding EM images in Fig.S1A, C and E, respectively. In Fig.S1G, the extent of the nanoclustering, *L*(*r*)-*r*, was plotted against the cluster radius, *r*, in nanometers for the corresponding EM images in Fig.S1A-F. The peak *L*(*r*)-*r* value, termed as *L*_*max*_, indicates the optimal nanoclustering (Fig.S1H). *L*_*max*_ value of 1 is the 99% confidence interval (99%C.I., green line), the values above which indicate statistically meaningful clustering. On the cell PM with further elevated curvature, the *L*_*max*_ of GFP-K-Ras^*G12V*^ or GFP-tK was markedly lower that the *L*_*max*_ values of GFP-H-Ras^*G12V*^ or GFP-tH (Fig.2D and Fig.S1J). When the cell PM was flattened, the nanoclustering of GFP-K-Ras^*G12V*^ or GFP-tK was significantly higher than that of the H-Ras constructs (Fig.2E and Fig.S1J). In the control experiments, the clustering of all Ras constructs was equivalent in untreated cells with normal curvature (Fig.2F and Fig.S1J). These spatial analyses consistently suggest that Ras proteins sense PM curvature modulations in an isoform-specific manner: K-Ras clustering favors less curved PM, whereas H-Ras clustering prefers more curved PM.

To better examine the extent of the PM curvature-induced changes in Ras oligomerization, we further interrogated our EM-spatial data and calculated the number of gold particles within a distance of 15nm to generate a population distribution. As shown in Fig.S2A-C, elevating the local PM curvature significantly increased the number of monomers but decreased the higher ordered multimers of GFP-K-Ras^*G12V*^. On the other hand, elevating PM curvature decreased monomer population and increased multimer population of GFP-tH (Fiug.S2D-F). To further validate the potential PM curvature-induced changes in K-Ras clustering/aggregation on the PM, we performed Raster image correlation spectroscopy combined with number / balance analysis (RICS-N/B) on live BHK cells. By counting the fluorescence intensity of GFP-K-Ras^*G12V*^, we estimated the population distribution of GFP-K-Ras^*G12V*^ on the PM of live BHK cells with normal PM curvature or with elevated PM curvature (expressing BAR_amph2_ domain). Further elevation of the PM curvature resulted in a marked shift in the population distribution of K-Ras^*G12V*^ oligomers on the PM (Fig.2G, Fig.S2G and H). Specifically, the presence of the BAR domain shifted the distribution of GFP-K-Ras^*G12V*^ towards monomers, when compared with the distribution of GFP-K-Ras^*G12V*^ in cells with normal curvature. To further examine whether the responses of Ras nanoclustering to changing PM curvature were dependent upon the specific BAR_amph2_ (an N-BAR) domain used above, we tested another BAR_FCC_ (an F-BAR) domain, which effectively induces PM curvature while sharing little sequence homology with the BAR_amph2_ domain ^25^. Fig.2H shows that the nanoclustering of GFP-K-Ras^*G12V*^ is markedly lower than that of GFP-tH in BHK cells expressing BAR_FCC_ (Elevated PM curvature), consistent with the effects of the BAR_amph2_ domain shown above. When the BAR_FCC_ is truncated to remove the coiled-coil domain, the resulting BAR_FCH_ domain still binds to the PM but no longer bends the membrane ^25^. In BHK cells expressing the BAR_FCH_ domain (nonBAR), the nanoclustering of both Ras isoforms did not respond (Fig.2H). Fig.2H also shows the control experiments, where the nanoclustering of GFP-K-Ras^*G12V*^ is similar to that of GFP-tH in untreated BHK cells with normal PM curvature. These BAR data, when combined together with the nanofabrication data, suggest that the response of the Ras nanoclustering to BAR domains is likely PM curvature-dependent.

To further validate the potential contribution of the cell PM to our observed changes in Ras clustering shown above, we next measured Ras oligomerization on the bilayers of the isolated native cell PM in the absence of all other intracellular constituents. We generated giant plasma membrane vesicles (GPMVs) from BHK cells. The GPMV is an ideal system for testing PM curvature sensitivity because they contain near-native PM constituents without any other cytoplasmic contents, such as cytoskeleton and endomembranes ^26^. The GPMVs containing GFP-tH with either empty vector pC1 or RFP-tH were subjected to HEPES buffers with different osmolality for 5 minutes before measuring the fluorescence lifetime of GFP, which was used to calculate FRET efficiency. The difference in osmolality across the bilayer is termed as ΔOsm, where increasing ΔOsm indicates gradual dilution of the external buffer, thus flattening of the bilayer. Fig.S2I shows that the FRET efficiency between GFP-tH and RFP-tH decreased as a function of elevating ΔOsm, suggesting that the oligomerization of the H-Ras anchor decreased dose-dependently as the vesicle membranes were flattened. Interestingly, the FRET efficiency between GFP- and RFP-tK showed a biphasic behavior: (1) below the ΔOsm threshold of ~100mM, the FRET efficiency increased as a function of elevating ΔOsm; (2) above the ΔOsm threshold of ~100mM, the FRET efficiency decreased as a function of elevating ΔOsm (Fig.S2I). The initial increase in FRET between GFP- and RFP-tK was likely caused by enhanced clustering upon flattening of the GPMV bilayer. The second FRET phase at high ΔOsm > 100mM was likely because the high hypotonic pressure over-stretched the bilayer and disrupted the global organization of the bilayer. Taken together, using multiple methods to manipulate membrane curvature in intact cell PM, live cells and isolated native PM, we show consistently that the spatiotemporal organization (PM localization and nanoclustering) of Ras proteins is dependent upon membrane curvature in an isoform specific manner: K-Ras (either full-length constitutively active oncogenic mutant K-Ras^*G12V*^ or the truncated minimal membrane-anchoring domain tK) favors less curved membranes, whereas H-Ras (either full-length oncogenic mutant H-Ras^*G12V*^ or the truncated membrane-anchoring tH) favors more curved membranes.

### Spatial segregation of PM lipids senses PM curvature in distinct manners

We next explored a potential mechanism mediating Ras membrane curvature sensing. In synthetic and *in silico* bilayers, the local distribution of lipids change at curved bilayer. Since Ras proteins undergo selective lipid sorting on the cell PM during the formation of signaling-essential nanoclusters, as membrane curvature sensing is potentially mediated via selective lipid sorting. However, the curvature-shifted lipid distribution in cells has been controversial ^21^. Specifically, how lipid headgroups and acyl chains influence their ability to sense membrane curvature in intact cells is not well established. To first quantify the spatial distribution of lipids upon PM curvature modulations in cells, we used EM-spatial analysis ^17,18^. PM lipid headgroups were probed via the GFP-tagged lipid-binding domains in BHK cells ^17,18^. The PM curvature was modulated as described above: hypotonic medium reduced PM curvature (Flat), untreated cell contained normal level of PM curvature (Normal), while cells expressing the BAR_amph2_ domain displayed further elevated PM curvature (Elevated) ^12,20,21,27,28^. The immunogold labeling analysis shows that the nanoclustering (Fig.3A) and gold labeling (Fig.3B) of GFP-LactC2, a specific PS probe, decreased as the PM curvature was gradually elevated. On the other hand, the nanoclustering (fig.3A) and the PM labeling (Fig.3B) of GFP-PH-PLCδ, a specific probe for PIP_2_, increased as the PM curvature elevated. GFP-D4H, a specific cholesterol probe, showed a similarly increased clustering (Fig.3A) and PM binding (Fig.3B) upon PM curvature elevation. The spatial distribution (Fig.3A) and PM binding (Fig.3B) of phosphoinositol 3,4,5-trisphosphate (PIP_3_) did not change upon PM curvature modulations. While the clustering of PA decreased as the cell PM was gradually elevated (Fig.3A), the dependence of PA levels in the PM on the PM curvature modulations was less clear (Fig.3B). Taken together, the lateral distribution and membrane localization of the anionic PM lipids respond to changing PM curvature in distinct manners.

**Figure 3.**
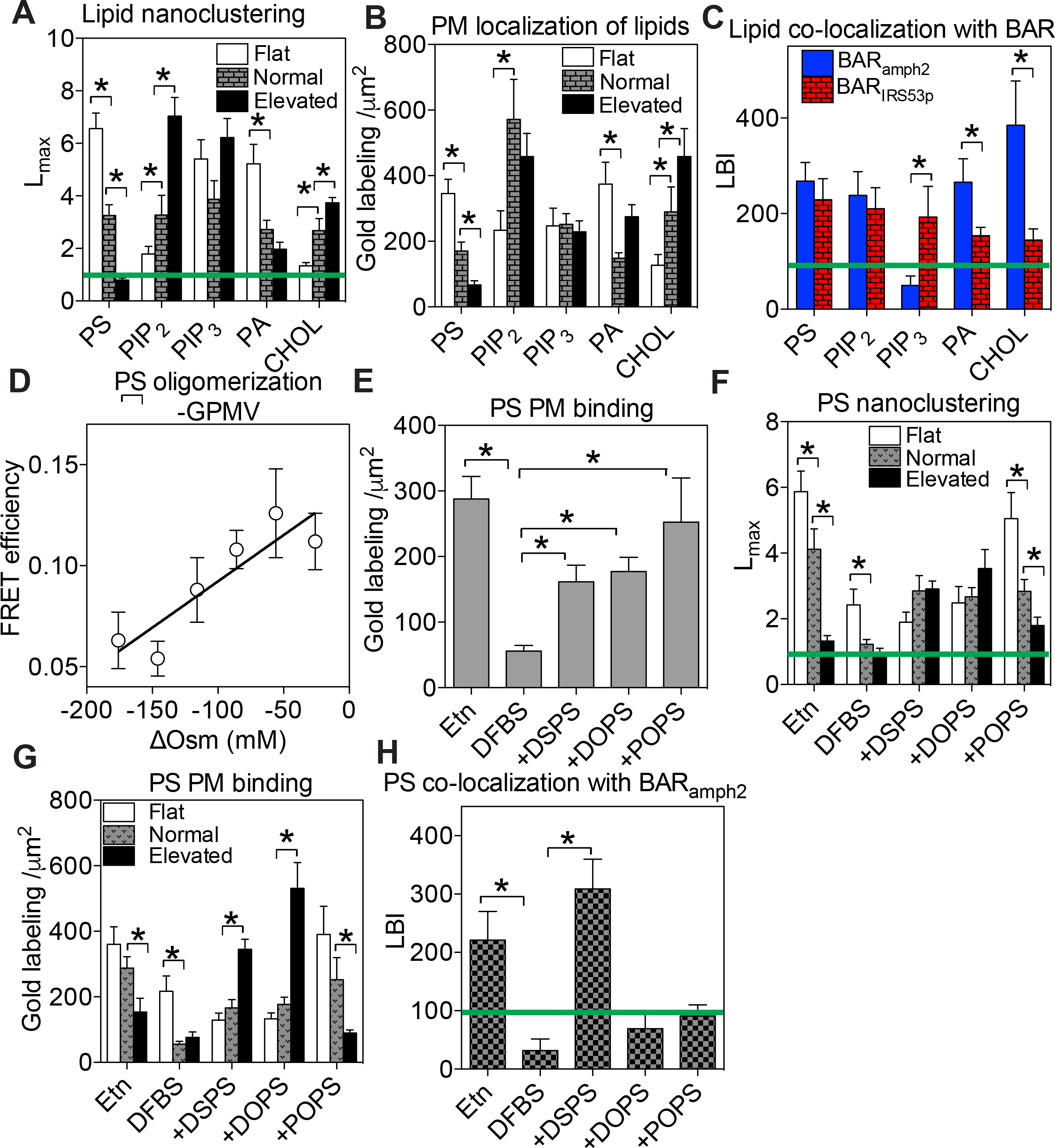
Plasma membrane lipids sense curvature modulations in distinct manners. (**A**) Optimal nanoclustering, *L*_*max*_, of GFP-tagged lipid-binding domains was calculated in BHK cells with flat PM (hypotonic incubation), normal PM curvature (untreated cells) or elevated PM curvature (ectopic co-expressing RFP-BAR_amph2_). The green line mark *L*_*max*_ value of 1 as the 99% C.I. (**B**) The number of gold nanoparticles within the 1μm^2^ area on the intact PM sheets was counted to estimate the localization of the GFP-tagged lipid-binding domains on the PM. (***C***) Co-localization, LBI, between the GFP-tagged lipid-binding domains and the RFP-tagged BAR domains was calculated. The LBI value of 100 indicates the 95%C.I. (marked by the green line), the values above which indicate statistically meaningful co-clustering. (**D**) The FRET efficiency between GFP- and RFP-tagged LactC2 on GPMVs was calculated using measurements of the fluorescence lifetime of GFP and was plotted as a function of increasing osmolality across the vesicle bilayers (ΔOsm). (**E**) The immunogold labeling of GFP-LactC2 on the PM is shown when PSA3 cells were under various endogenous PS manipulations: Etn →recovery of the normal endogenous PS; DFBS → endogenous PS depletion; +DSPS → addback of di18:0 PS under DFBS condition; +DOPS → addback of di18:1 PS under DFBS; and +POPS → addback of 16:0/18:1 PS under DDFBS condtion. *L*_*max*_ (**F**) and PM gold labeling (**G**) of GFP-LactC2 in PSA3 cells under various PS-manipulating and PM curvature modulation conditions was calculated. (**H**) Co-localization, LBI, between the GFP-LactC2 and the RFP-BAR_amph2_ was calculated. All values are shown as mean±SEM. The statistical significance in the univariate and bivariate spatial analyses (*A*, *C*, *F*, *G* and *H*) was evaluated using the bootstrap tests with * indicating p < 0.05. The statistical significance of the gold labeling (*B*, *E* and *G*), as well as the FLIM-FRET data (*D*), was evaluated using the one-way ANOVA with * indicating p < 0.05.

To further characterize potential curvature preference of various lipids in the PM, we quantified co-localization between the GFP-tagged lipid-binding domains and the RFP-tagged BAR domains inducing opposite curvature directions: RFP-BAR_amph2_ to induce negative curvature or RFP-BAR_IRS53p_ to induce positive curvature. GFP and RFP on the intact PM sheets attached to the EM grids were immunolabeled with 6nm gold conjugated to anti-GFP antibody and 2nm gold conjugated to anti-RFP antibody, respectively. The gold nanoparticles were imaged using TEM and the spatial co-localization between the two populations of the gold particles was calculated using the Ripley’s bivariate K-function analysis (Fig.S3A and B). The extent of the co-clustering, *L*_*biv*_(*r*)-*r*, was plotted against the cluster radius, *r*, in nanometers (Fig.S3C). To summarize the co-localization data, we calculated the area-under-the-curve values for all the *L*_*biv*_(*r*)-*r* curves between the cluster radii (*r*) range of 10-110nm to yield values of L-function-bivariate-integrated (LBI, Fig.S3C). The higher LBI values indicate more extensive co-localization, with an LBI value of 100 as the 95% confidence interval (95%C.I., the green line in Fig.S3C). PS, PIP_2_, PA and cholesterol, but not PIP_3_, co-localized with BAR_amph2_. All lipids tested co-localized with BAR_IRS53p_. Further, PA and cholesterol co-localized more extensively with BAR_amph2_ than with BAR_IRS53p_ (Fig.3C). Thus, the spatial packing of PM lipids responds to curvature modulations in distinct manners.

To test the curvature preference of lipid tails in intact cells, we focused on the most abundant anionic phospholipid, PS, using the PS auxotroph PSA3 cells. With a PS synthase (PSS1) knocked down, the PSA3 cells generate significantly less endogenous PS when grown in dialyzed fetal bovine serum (DFBS) ^17,18,29^. Supplementation of 10μM ethanolamine (Etn) stimulates PSS2 and effectively recovers the wild-type level of the endogenous PS level ^17,18,29^. Immunogold-labeling of GFP-LactC2 confirms the changes in PS levels in the PM inner leaflet (Fig.3E). Addback of different synthetic PS species to the PSA3 cells under the condition of endogenous PS depletion (DFBS) resulted in the selective enrichment of distinct PS species, which has been verified in lipidomics ^17^ (Fig.3E). The nanoclustering (Fig.3F) and PM binding (Fig.3G) of 16:0/18:1PS (POPS) were gradually disrupted upon elevating PM curvature. On the other hand, the clustering of the fully saturated di18:0 PS (DSPS) and the mono-unsaturated di18:1 PS (DOPS) did not respond to changing PM curvature (Fig.3F). Interestingly, the immunogold labeling of GFP-LactC2 increased markedly upon elevating PM curvature in the presence of either DSPS or DOPS (Fig.3G), suggesting that the PM localization of both PS species favors more curved PM although their spatial segregation did not change. We next tested co-localization of different PS species with the BAR_amph2_-induced PM curvature. Only the fully saturated DSPS co-localized extensively with the BAR_amph2_ domain (Fig.3H). Thus, PS acyl chains are sensitive to changing PM curvature.

### Distinct PS species mediate K-Ras membrane curvature sensing

K-Ras nanoclusters contain distinct PS species, specifically mixed-chain PS species ^17^. Further, effector recruitment by the constitutively active oncogenic mutant K-Ras^*G12V*^ occurs only in the presence of the mixed-chain PS but not other PS species tested ^17^ We, here, show that the nanoclustering, PM binding and PM curvature affinity of different PS species responded to changing PM curvature in distinct manners (Fig.3E-H). To test if different PS species selectively mediated K-Ras PM curvature sensing, we used surface plasmon resonance (SPR) to measure the binding of the purified full-length GTP-bound K-Ras to large unilamellar vesicles (LUVs) of different sizes, an established method to manipulate bilayer curvature ^14^. On LUVs composed of POPS/POPC (20:80), the binding of K-Ras increased as the diameters of the LUVs became larger (Fig.4A), suggesting that K-Ras binding to the bilayers favors less curved membranes. On LUVs of DOPS/DOPC (20:80), K-Ras binding was independent of vesicle size (Fig.4B). These data clearly suggest that different PS species selectively mediate the membrane curvature-dependent binding of K-Ras. We then tested the PS selectivity on intact cell PM. The endogenous PS depletion in PSA3 cells (DFBS) effectively abolished the changes in the nanoclustering of GFP-K-Ras^*G12V*^ upon hypotonic flattening of the PM (Fig.4C). Wild-type PS level (Etn) restored the PM curvature sensitivity of GFP-K-Ras^*G12V*^ (Fig.4C). We then tested the potential selectivity of various PS species in cells via supplementation of synthetic PS species under the endogenous PS depletion (DFBS). Addback of only POPS, but not DSPS or DOPS, effectively restored the curvature sensitivity of GFP-K-Ras^*G12V*^ (Fig.4C). Thus, different PS species selectively mediate K-Ras PM curvature sensing consistently in intact cells and synthetic vesicles.

**Figure 4.**
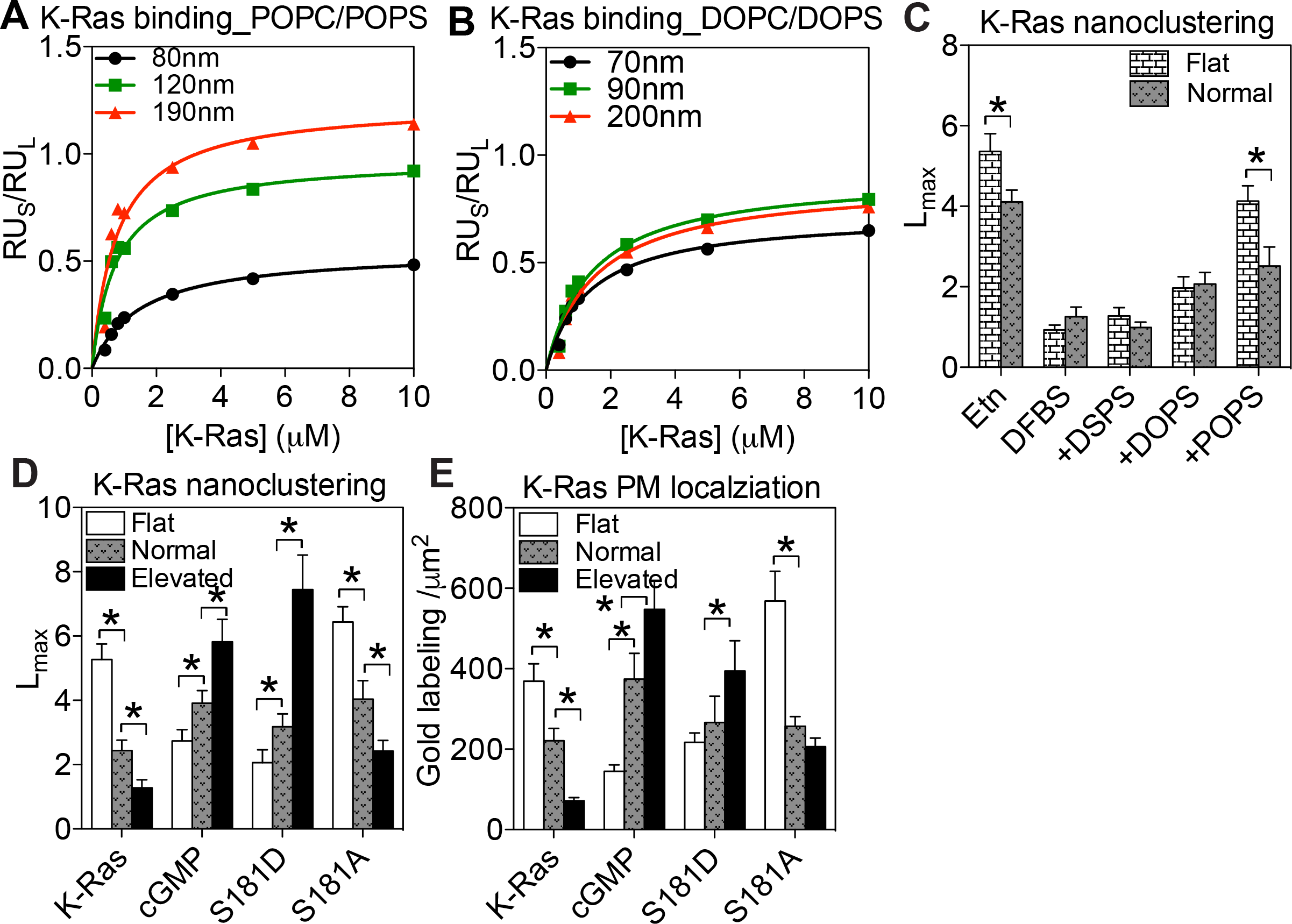
PS mediates K-Ras PM curvature sensing. SPR was used to measure binding of the purified full-length K-Ras to different-sized LUVs composed of POPS/POPC (20:80) (**A**) or DOPS/DOPC (20:80) (**B**). (**C**) PSA3 cells expressing GFP-K-Ras^*G12V*^ were grown under various PS-manipulation conditions. *L*_*max*_ of GFP-K-Ras^*G12V*^ was analyzed at isotonic vs. hypotonic conditions. *L*_*max*_ (**D**) and PM binding (**E**) of GFP-tagged phosphorylation mutants of K-Ras^*G12V*^ in BHK cells were quantified upon PM curvature manipulation. All data are shown as mean±SEM. The statistical significance in the clustering analyses was evaluated using the bootstrap tests, with * indicating p<0.05. The statistical significance of the gold labeling was evaluated using the one-way ANOVA with * indicating p<0.05.

To further examine whether K-Ras PM curvature sensing is mediated by lipid sorting, we compared the clustering of unphosphorylated vs. phosphorylated K-Ras^*G12V*^ in BHK cells. Because of the K-Ras phosphorylation site Ser181 is located in its minimal membrane anchoring domain immediately adjacent to its polybasic domain (Lysines 175-180), Ser181 phosphorylation switches K-Ras lipid preference from PS to PIP_2_ ^17^ without interfering with its ability to recruit downstream effectors ^30,31^. As PIP_2_ local segregation favors more curved membranes, opposite of PS (Fig.3A and B), comparing unphosphorylated vs. phosphorylated K-Ras^*G12V*^ is ideal for examining the potential lipid dependence of K-Ras PM curvature sensing. The nanoclustering and PM binding of GFP-K-Ras^*G12V*^ in untreated cells, or an unphosphorylatable mutant GFP-K-Ras^*G12V/S181A*^, responded consistently (Fig.4D and E). On the other hand, the clustering and PM binding of the phosphorylated GFP-K-Ras^*G12V*^ in BHK cells treated with 500µM cGMP ^17^ or those of a phospho-mimetic mutant GFP-K-Ras^*G12V/S181D*^ were gradually enhanced upon PM curvature elevation (Fig.4D and E). Thus, the phosphorylated K-Ras switches its PM curvature preference to more curved PM. Taken together, K-Ras membrane curvature sensing is largely driven by its ability to selectively sort lipids in the PM.

### K-Ras-dependent MAPK signaling is sensitive to PM curvature modulations

We then tested Ras signaling upon changing PM curvature. Pre-serum-starved wild-type BHK cells were subjected to media with various ΔOsm for 5 minutes. Increasing ΔOsm across the cell PM gradually flattens the PM. Levels of phosphorylated MEK and ERK in the K-Ras-regulated MAPK cascade and pAkt in the H-Ras-regulated PI3K cascade were evaluated using Western blotting. Increasing ΔOsm elevated the pMEK/pERK, but decreased the pAkt (Fig.5A and Fig.S4A). To test reversibility, BHK cells were incubated in the hypotonic medium for 5 minutes before switching back to the isotonic medium for various time points. The MAPK and the PI3K signaling recovered to the baseline between 15 and 30 minutes following switch back to the isotonic medium (Fig.5B and Fig.S4B). To further validated the role of K-Ras in the MAPK response to changing PM curvature, we used various Ras-less mouse epithelial fibroblast (MEF) lines, which have all the endogenous Ras isoforms knocked down. We focused specifically on 3 MEF lines: (1) K-Ras^*G12V*^ MEF line contains only the constitutively active oncogenic mutant K-Ras^*G12V*^ but no other endogenous Ras isoforms, (2) BRaf^*V600E*^ MEF line contains the constitutively active oncogenic mutant of a K-Ras effector BRaf^*V600E*^ and all the endogenous Ras isoforms have been knocked down, (3) wild-type MEF line contains all the endogenous Ras isoforms ^32^. The pERK level in the K-Ras^*G12V*^-MEF increased upon hypotonic condition more efficiently than that in the wild-type MEF line containing endogenous Ras isoforms (Fig.4C). The MAPK signaling in the BRAF^*V600E*^-MEF cells (no endogenous Ras isoforms) no longer responded to the osmotic imbalance. Thus, K-Ras likely drives MAPK response to PM curvature manipulations.

**Figure 5.**
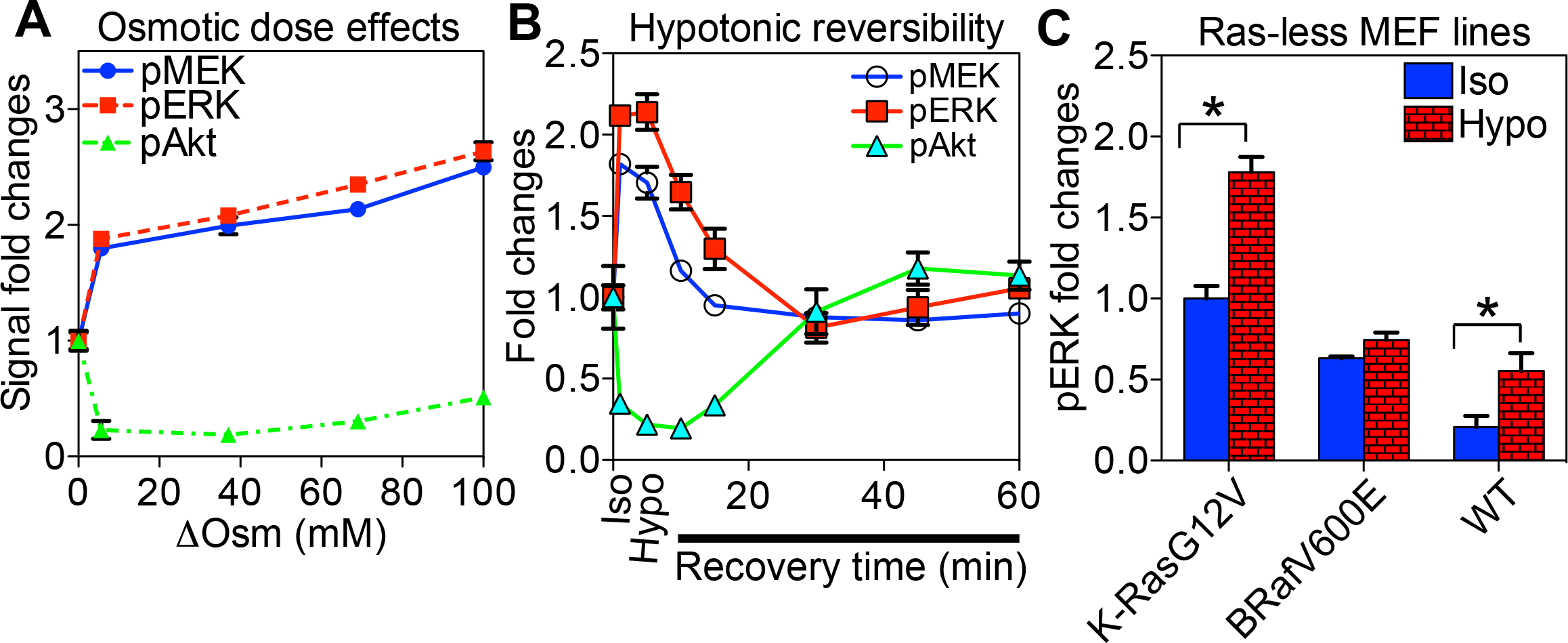
Ras signaling senses PM curvature modulations in an isoform-specific manner. (**A**) BHK cells were pre-serum-starved for 2h and subjected to the media at various osmolality values for 5 minutes before harvesting. Levels of the phosphorylated MAPK components, pMEK and pERK, were quantified using Western blotting. (**B**) Pre-serum-starved BHK cells were incubated in the media pre-diluted with 40% water for 5 minutes before being returned to the isotonic media for various time periods. (**C**) Ras-less MEF lines stably expressing K-Ras^*G12V*^, BRAF^*V600E*^ or wild-type MEF were subjected to isotonic or hypotonic (diluted with 40% H_2_O) HEPES buffers for 5 minutes before harvesting. For each experiment, the whole cell lysates were used for blotting against pMEK, pERK and pAkt. Quantitation of 3 independent experiments is shown as mean±SEM. All values were normalized against the perspective bands in untreated cells in isotonic medium. The statistical significance was evaluated using the one-way ANOVA, with * indicating statistical significance (p<0.05).

## Discussion

Mitogenic signaling that regulate cell growth, division, proliferation and migration, correlates tightly with changing cell morphology. How fluctuations in cell PM curvature during changing cell shape are correlated with the intracellular MAPK signaling is still poorly understood. The lipid-anchored Ras small GTPases are the upstream regulators of the MAPK/PI3K cascades and undergo selective lipid sorting on the cell PM for efficient signal transduction. Thus, Ras function may be regulated by the changing PM curvature. Indeed, the membrane partitioning of the truncated minimal membrane anchoring domains of H-Ras and N-Ras displays curvature sensitivity in synthetic bilayers or *in silico* ^14,15,33^. In our current study, we used various methods to manipulate PM curvature of intact/live cells and compared the signaling-essential spatiotemporal organization of the full-length constitutively active oncogenic Ras mutants, K-Ras^*G12V*^ and H-Ras^*G12V*^, as well as their truncated minimal membrane anchors tK and tH.

Our findings on the PM curvature preferences of the H-Ras minimal anchor in intact/live cells are consistent with the predictions from the MD simulations, which suggest that tH preferentially localizes to the boundaries between liquid-ordered (*L*_*o*_) and liquid-disordered (*L*_*d*_) domains and induce bilayer bending ^33^. This makes sense because tH contains two fully-saturated palmitoyl chains and a poly-unsaturated farnesyl chain, which favor domain boundaries. Similarly, the localization of the N-Ras anchor tN, which also possesses a palmitoyl chain, favorably partitions into the *L*_*o*_ domains of the highly curved model bilayers ^14,15^. K-Ras membrane curvature sensing has not been studied before. The membrane curvature preference of H- and N-Ras cannot be extrapolated to K-Ras. Although the three Ras isoforms share near identical enzymatic G-domains, Ras functions have been consistently shown to be isoform-specific in patients, as well as model systems such as animal models, cultured cells, synthetic bilayers, and MD simulations ^17–19,34–45^. Specifically, knocking down only K-Ras, but not other Ras isoforms, is embryonically lethal ^43–45^. Ras oncogenic mutations exist in distinct tissues: K-Ras in pancreas, colon and lungs; N-Ras in skin and myeloma; and H-Ras in head and neck ^4,5^. K-Ras is the most prevalent isoform in cancer and its oncogenic mutations contribute to ~80% of all Ras-related cancers ^4^. Thus, studying K-Ras presents a unique set of challenges not found in other Ras isoforms. Because of the unique C-terminal membrane-anchoring farnesylated hexa-lysine polybasic domain not shared by other Ras isoforms, K-Ras sorts distinct set of PM lipids than other Ras isoforms to form nanoclusters, which results in the isoform-specific effector recruitment and signal propagation ^16–19,24,36,46^. Here, we report that the spatiotemporal organization of K-Ras (either the full-length oncogenic mutant K-Ras^*G12V*^ or the truncated minimal anchor tK) favors less curved membranes than that of H-Ras in intact/live cells, as well as isolated native PM vesicles. The potential contribution of K-Ras in the cell shape-dependence of MAPK signaling was further validated in signaling experiments using a series of Ras-less MEF lines. Strikingly, the MAPK signaling in the MEF line expressing only the oncogenic mutant K-Ras^*G12V*^ (but none of the endogenous wild-type Ras isoforms) is more sensitive to hypotonicity-induced cell shape changes than the wild-type MEF expressing all endogenous wild-type Ras isoforms. Further, the MEF line expressing the oncogenic mutant of the K-Ras downstream effector BRAF^*V600E*^ (but none of the Ras isoforms) becomes insensitive to changing cell shape. These signaling data, together with the spatial analysis, suggest that K-Ras functions, including those of the oncogenic K-Ras mutants, are regulated by PM curvature modulations during cell shape changes.

We then focused on K-Ras for exploring a potential mechanism for its membrane curvature sensing because of its important activities in cancer. Because we have shown previously that K-Ras selective sorts PS in nanocustering, effector binding and signaling, we propose that PS is an important factor mediating K-Ras curvature sensing. The potential roles of lipids in mediating the membrane curvature sensing of proteins have been controversial. Recent studies suggest that lipids, even those with strong spontaneous curvatures such as PIP_2_, only play minor roles in mediating protein curvature sensing ^47^. However, the lipid-anchored small GTPases rely heavily on merely fatty acid chains for membrane anchoring, with their globular G-domains mostly hanging off the membrane. In the case of K-Ras, several lines of evidence in our current study suggest that the selective interactions between K-Ras and mixed-chain PS species may contribute to K-Ras membrane curvature sensing. (1) Our EM data show that the spatiotemporal organization of either the full-length K-Ras^*G12V*^ or the truncated minimal membrane-anchoring domain tK respond to changing PM curvature in a similar manner. In the absence of the G-domain (amino acids 1-166), tK only contains a short hexa-lysine polybasic domain (amino acids 175-180) with an adjacent poly-unsaturated farnesyl chain conjugated to Cys185. Thus, the lateral distribution of tK on the PM is mostly driven by the biophysical interactions between tK and PM lipids. (2) Our *in vitro* assay clearly demonstrates that the binding of the purified K-Ras to the synthetic LUVs depends on the sizes of the vesicles only in the presence of particular PS species, suggesting that K-Ras bilayer curvature preferences depend on PS acyl chain structures and pointing to the important roles of PS in mediating K-Ras curvature sensing. (3) The spatial segregation of only mixed-chain PS species, but not other PS species tested, respond to changing PM curvature in a similar manner as that of K-Ras, supporting the idea that PS mediates K-Ras PM curvature sensing. (4) In intact cells, K-Ras PM curvature sensing is abolished when the endogenous PS is depleted, and is recovered upon addback of the mixed-chain PS species but not other PS species tested. This data further showcases the important roles of PS in mediating K-Ras PM curvature sensing. (5) The spatiotemporal organization of the phosphorylated K-Ras, which switches its lipid selectivity to PIP_2_ instead of PS ^17^, favors highly curved PM. This is the opposite of the PS-sorting unphosphorylated K-Ras, but similar to the behavior of PIP_2_. As the K-Ras phosphorylation site Ser181 is located within the minimal membrane anchoring domain (amino acids 175-185), the phosphorylation has no effect on the enzymatic activities of K-Ras and only alters how K-Ras localizes to the PM ^30,31^ and the selective lipid sorting ^17^. Taken together, these data consistently point to the possibility that distinct PS species selectively mediate K-Ras membrane curvature sensing.

Although specific interactions between membrane proteins and BAR domains may contribute to their membrane curvature sensing, it is unlikely that K-Ras uses this mechanism in its PM curvature sensing. While we used ectopically expressed BAR domains to further elevate PM curvature, we also manipulated PM curvature using many methods in the absence of over-expressed BAR domains. We observed consistent responses from either the full-length K-Ras^*G12V*^ or the minimal anchoring domain tK to BAR-induced membrane curvature modulations. Thus, it is unlikely Ras membrane curvature sensing is caused by specific interactions between Ras and the BAR domains. Rather, selective sorting of PM lipids by Ras is likely a mechanism mediating Ras curvature sensing.

An interesting observation in our current study is that different PS species with distinct acyl chain structures respond to changing PM curvature in distinct manners. Electrostatic interactions have been thought to dominate the interactions between the curvature-inducing BAR domains enriched with positively charged residues and the anionic lipid headgroups. The different PS species with the same headgroup but distinct acyl chain compositions should associate with the BAR domain in similar manner. Yet, we observed distinct responses by different PS species to changing PM curvature. Different PS species also co-localize with the BAR domain in distinct manners, suggesting that more than electrostatic interactions between the BAR domain and the lipid headgroups contribute to the observed selectivity. Our observed curvature preferences of different PS species are consistent with the distinct spontaneous curvature of these PS species. Specifically, we show that the univariate spatial distribution of mixed-chain PS favors less curved PM, whereas the clustering of symmetric PS species favors more curved membranes in intact cells. Bivariate co-localization analyses further show that only fully saturated PS, but not other PS species, co-localizes with the inwardly bending BAR_amph2_ domain. The spontaneous curvature of PS species is not well-established. We can use PC species as reference because both PS and PC are cylindrical lipids and possess similar packing geometries. The fully saturated PC species, such as di14:0 PC (DMPC) and di16:0 PC (DPPC), possess positive spontaneous curvature (*J_o_*) values. Lipids with positive *J_o_* values prefer BAR-induced curved bilayers ^48^. On the other hand, unsaturated PC species, such as di18:1 PC (DOPC), have negative *J_o_* values (−0.09 nm^-1^) and likely does not fit well with the N-BAR-induced inwardly bent membranes. Mixed-chain PC species, such as 16:0/18:1 PC (POPC), have the lowest *J_o_* values (−0.022 nm^-1^), thus less likely to pack well in curved membranes. This is consistent with our observations to suggesting that the PM curvature preferences of different PS species correlate with their packing geometries.

## Materials and Methods

### Materials

GFP-BAR_amph2_, GFP-BAR_EFC_ and GFP-BAR_FCH_ have been generously provided by Dr. Pietro De Camilli at the Yale University.

### Methods

#### Electron microscopy (EM)-spatial analysis

##### EM-univariate spatial analysis

The univariate K-function analysis calculates the nanoclustering of a single population of gold immunolabeling on intact PM sheets ^17,18^. Intact PM sheets of cells ectopically expressing GFP-tagged proteins/peptides of interest were attached to copper EM grids. After fixation with 4% paraformaldehyde (PFA) and 0.1% gluaraldehyde, GFP on the PM sheets was immunolabeled with 4.5nm gold nanoparticles conjugated to anti-GFP antibody and negative-stained with uranyl acetate. Gold distribution on the PM was imaged using TEM at 100,000x magnification. The coordinates of every gold particle were assigned via ImageJ. Nanoclustering of gold particles within a selected 1μm^2^ area on intact PM sheets was quantified using Ripley’s K-function. The test is designed to test a null hypothesis that all points in a selected area are distributed randomly:

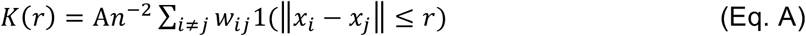

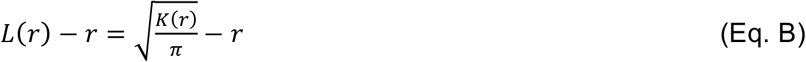

where *K*(*r*) indicates the univariate K-function for *n* gold nanoparticles in an intact PM area of *A*; *r* is the length scale between 1 and 240 nm with an increment of 1nm; ‖ · ‖ is Euclidean distance; where the indicator function of 1(·) = 1 if ‖*x*_*i*_-*x*_*j*_‖ ≤ r and 1(·) = 0 if ‖*x*_*i*_-*x*_*j*_‖ > r. To achieve an unbiased edge correction, a parameter of *w*_*ij*_^−1^ is used to describe the proportion of the circumference of a circle that has the center at *x*_*i*_ and radius ‖*x*_*i*_-*x*_*j*_‖. *K*(*r*) is then linearly transformed into *L*(*r*) − *r*, which is normalized against the 99% confidence interval (99% C.I.) estimated from Monte Carlo simulations. A *L*(*r*) - *r* value of 0 for all values of *r* indicates a complete random distribution of gold. A *L*(*r*) - *r* value above the 99% C.I. of 1 at the corresponding value of *r* indicates statistical clustering at certain length scale. At least 15 PM sheets were imaged, analyzed and pooled for each condition in the current study. Statistical significance was evaluated via comparing our calculated point patterns against 1000 bootstrap samples in bootstrap tests ^17,18^.

##### EM-Bivariate co-localization analysis

Co-localization between two populations of gold immunolabeling GFP-tagged and RFP-tagged proteins / peptides is quantified using the bivariate K-function co-localization analysis ^17,18^. Intact apical PM sheets of cells co-expressing GFP- and RFP-tagged proteins / peptides were attached and fixed to EM grids. The PM sheets were immunolabeled with 2nm gold conjugated to anti-RFP antibody and 6nm gold linked to anti-GFP antibody, respectively. X / Y coordinates of each gold nanoparticle were assigned in ImageJ and the co-localization between the two gold populations was calculated using a bivariate K-function. The analysis is designed to test the null hypothesis that the two point populations spatially segregate from each other. (Eqs. C-F):

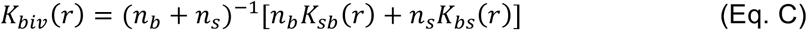

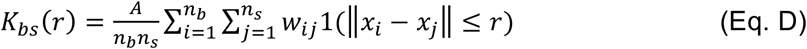

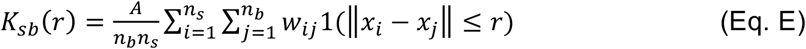

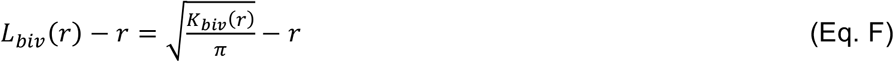

where *K*_*biv*_(*r*) is the bivariate estimator comprised of 2 separate bivariate K-functions: *K_bs_*(*r*) describes the distribution of all the big 6nm gold particles (*b* = big gold) with respect to each 2nm-small gold particle (*s* = small gold); and *K_sb_*(*r*) describes the distribution of all the small gold particles with respect to each big gold particle. The value of n_b_ is the number of 6nm big gold particles and the value of n_s_ is the number of 2nm small gold particles within a PM area of *A*. Other notations follow the same description as explained in Eqs.A and B. *K_biv_*(*r*) is then linearly transformed into *L*_*biv*_(*r*)-*r*, which was normalized against the 95% confidence interval (95% C.I.). An *L*_*biv*_(*r*)-*r* value of 0 indicates spatial segregation between the two populations of gold particles, whereas an *L*_*biv*_(*r*)-*r* value above the 95% C.I. of 1 at the corresponding distance of *r* indicates yields statistically significant co-localization at certain distance yields. Area-under-the-curve for each *L*_*biv*_(*r*)-*r* curves was calculated within a fixed range 10 < *r* < 110 nm, and was termed bivariate *L*_*biv*_(*r*)-*r* integrated (or LBI):

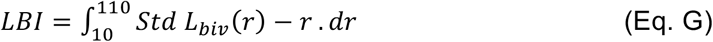

For each condition, > 15 apical PM sheets were imaged, analyzed and pooled, shown as mean of LBI values ± SEM. Statistical significance between conditions was evaluated via comparing against 1000 bootstrap samples as described ^17,18^.

#### FLIM-FRET in GPMVs

BHK cells co-expressing GFP-tagged Ras anchors and the empty vector pC1 or RFP-tagged Ras anchors were grown to ~85% confluency and washed with 2x HEPES buffer. Cells were then incubated in HEPES buffer containing 2mM NEM for ~90 minutes to induce blebbing ^49^. GPMVs containing GFP- and RFP-tagged Ras anchors (or GFP-tagged Ras anchors with pC1) were then incubated in HEPES buffers containing various percentages of de-ionized water (hypotonic conditions) or concentrations of NaCl (hypertonic conditions) for 5 minutes.

GFP fluorescence was visualized using a Nikon TiE wide-field microscope using a 60x oil-emersion PLAN-Apo / 1.4 numerical aperture lens ^17,18^. The fluorescence lifetime of GFP was measured using a Lambert FLIM unit attached to the wide-field microscope. GFP was excited using a sinusoidally stimulated and modulating 3-watt 497nm light-emitting diode (LED) at 40Hz. At least 20 vesicles were imaged and the fluorescence lifetime values were pooled and averaged. Statistical significance was evaluated using one-way ANOVA, with * indicating p < 0.05.

#### Fabrication and Characterization of Quartz Nanostructures

##### Nano-fabrication

Nanostructures used in this work were fabricated on the square quartz wafer by using electron-beam lithography. In brief, the chips were spin-coated with 300nm of positive electron-beam resist PMMA (MicroChem), followed by AR-PC 5090.02 (Allresist). Desired patterns were exposed by EBL (FEI Helios NanoLab), then, developed in IPA:MIBK=3:1 solution. A 100nm Cr mask was generated via thermal evaporation (UNIVEX 250 Benchtop) and lift-off in aceton. Nanostructures were subsequently synthesized through reactive ion etching with the mixture of CF_4_ and CHF_3_ (Oxford Plasmalab80). Prior to cell culture, the nanostructured chips were cleaned by O_2_ plasma and immersed in Chromium Etchant (Sigma-Aldrich) to remove the remaining Cr mask. SEM (FEI Helios NanoLab) imaging was performed after 10 nm-gold-coating to measure the dimensional properties of different nanostructures.

##### Cell culture

The nanostructure substrates were firstly coated with fibronectin (2ug/ml, Sigma-Aldrich) for 30mins at 37°C. After coating, U2OS cell culture was plated onto the substrates with 400000 cells per 35mm dish one night before transfection. The cells were maintained in the DMEM supplemented with 10% fetal bovine serum (FBS) (Life Technologies) and 1% Penicillin-Streptomycin (Life Technologies) in a standard incubator at 37°C with 5% CO_2_ overnight.

##### Transfection

For GFP-K-RasG12V and GFP-tH transfection in U2OS cells, 1 μg plasmid was mixed with 1.5 μl Lipofectamine 3000 (Life Technologies) and 2 μl P3000 reagent (Life Technologies) in Opti-MEM (Gibco) and incubated for 20 mins at room temperature. The medium of the cells to be transfected was then changed to Opti-MEM and the transfection mixture was added. Next, the cells were incubated at 37°C in Opti-MEM. After 4 hours, the medium was changed back to regular culture medium and the cells were allowed to recover for additional 4 hours before imaging.

##### Live-cell imaging and quantification

Live imaging of the transfected U2OS cells on regular nanobar arrays (250 nm wide and 2 μm long) was performed using laser scanning confocal microscopy (Zeiss LSM 800 with Airyscan). In particular, a Plan-Apochromat 100x/1.4 oil objective was used. Excitation of EGFP was performed at 488nm and detection was at 400- 575nm. Images had a resolution of 1024×1024 pixels, with a pixel size of 62nm and a bit depth of 16. During imaging, cells were maintained at 37°C in an on-stage incubator with FluoroBrite DMEM (Gibco). Z stack images were acquired at 512×512 pixels with 100nm distance between frames.

The curvature preferences of KG12V and tH (Fig.2) were measured on regular nanobar arrays. After averaging the z-stack images captured using confocal, the background intensity of each image was subtracted by a rolling ball algorithm in Fiji with 3-pixel radius. The protein intensities at nanobar ends and centres were quantified using a custom-written Matlab (MathWorks) code and bar-end / bar-center ratio were calculated and displayed as mean ± s.e.m as indicated in the figure.

#### Raster image correlation spectroscopy-number and balance analysis (RICS-NnB)

Fluorescence measurements for N & B analysis were carried out with Nikon A1 confocal microscope using CFI Plan Apo IR 60× 1.27 NA water immersion objective at 22 °C. Live cells were maintained at 37 °C in Dulbecco’s modified Eagle medium (DMEM), 10% bovine calf serum (Hyclone) and 5% (v/v) CO_2_ in a 35 mm glass bottom dish (MatTek Corporation). Cells were imaged in live cell imaging solution containing HEPES buffered physiological saline at pH 7.4 (Life Technologies). Fluorescence images were acquired using a 488 nm laser power at 0.5% that corresponded to 1.75 mW at the measured samples. Rest of the measuring parameters and microscope settings were similar to what has been described in previous report 50. Briefly, for the N & B analysis, acquired images were scanned with 64 × 64 pixels box along the cell margins. Brightness values of the BHK cells expressing mem-EGFP (membrane-bound EGFP, a GFP containing an N terminally palmitoylated GAP-43 sequence cloned in a pEGFP-1 vector, have been used as brightness standards and for calibrating laser power and other microscope parameters. Our data have been analyzed following previous reports ^50^. Oligomeric size of KRAS has been determined as the ratio of the measured brightness and the brightness of monomeric mem-EGFP after subtracting the apparent brightness of immobile molecules.

#### Surface plasmon resonance (SPR)

Fully processed KRAS4b was produced and purified as described before ^51^. All lipids were purchased from Avanti Polar Lipids (Alabaster, USA): 1,2-dioleoyl-*sn*-glycero-3 phosphocholine (DOPC), 1,2-dioleoyl-sn-glycero-3-phospho-L-serine sodium salt (DOPS), 1-palmitoyl-2-oleoyl-*sn*-glycero-3-phosphocholine (POPC), 1-palmitoyl-2-oleoyl-*sn*-glycero-3-phospho-L-serine sodium salt (POPS).

##### Liposome preparation

Lipid stock solutions dissolved in chloroform were mixed at the desired molar composition. Chloroform was evaporated in a gaseous nitrogen stream and afterward dried overnight under vacuum to remove the residual solvent. Before the experiments, the lipids were suspended in ~1 mL 20 mM HEPES buffer (pH 7.4), 150 mM NaCl, 1 mM TCEP and vortexed to yield a theoretical total lipid concentration of 6 mM. After hydration, the lipid solution was sonicated for 5 min at 25 °C, and subjected to five freeze-thaw-vortex cycles and another brief sonication. Large unilamellar vesicles (LUVs) were formed by extrusion through a polycarbonate filter ranging from 30 to 400-nm pore diameter. The extruded solution was then diluted to a concentration of 1 mM.

##### Surface Plasmon Resonance

SPR experiments were carried out in a Biacore S200 instrument from GE Healthcare. Temperature was set at 25°C for all experiments. A 20 mM HEPES, 150 mM NaCl, pH 7.4, 5 μM GppNHp solution was used as running buffer for experiments using K-Ras in the active state. The flow system was primed three times before initiating an experiment. The L1 sensor chip was used in all experiments. The sensor chip surface was rinsed with three injections of 20 mM CHAPS before LUV deposition. 1 mM lipid LUV samples were injected over the L1 sensor chip for 900 s, at a 2 μL/min flow rate. Loose vesicles were removed with a 36 s injection of 10 mM NaOH at 50 μL/min. K-Ras at defined concentrations (0.2 - 50 μM) were injected over pre-formed lipid vesicle-coated surfaces at 30 μL/min, for a total of 60 s (association phase). Solutes were allowed to dissociate for 300 s. L1 sensor chip surface regeneration was performed with sequential injections of 20 mM CHAPS (5 μL/min for 60 s). Baseline response values were compared before and after each experiment to evaluate the effectiveness of the surface regeneration. Raw SPR sensorgram data were collected for both lipid deposition and solute binding. LUV deposition response values were collected from sensorgrams upon reaching a stable response. For each studied molecule, association steady-state response values were collected from individual sensorgrams at *t* = 60 s. Dissociation response data were collected between 60 and 360 s of each sensorgram.

#### Western blotting

BHK cells were grown in DMEM containing 10% BCS to ~85-90% confluency and pre-serum-starved for 2h before incubation with hypotonic medium containing various percentages of de-ionized water for 5 minutes. Whole cell lysates were collected. In reversibility experiments, pre-serum-starved BHK cells were exposed to hypotonic medium containing 40% water for 5 minutes before being returned to isotonic medium for various time length before harvesting. Whole cell lysates were used and blotted using antibodies against pMEK, pERK and pAkt. Actin was used as loading control. Each experiment was performed at least 3 times separately and data was shown as mean±SEM. Statistical significance was evaluated using one-way ANOVA, with * indicating p < 0.05.

## Author contributions

WZ, AAG and YZ designed experiments; HL, HM, SSB, BL, FJF, WZ and YZ performed experiments; WZ, DVN, AAG and YZ wrote manuscript.

## Data and materials availability

All data is available in the main text or the supplementary materials.

---

**Supplemental Figure S1.**
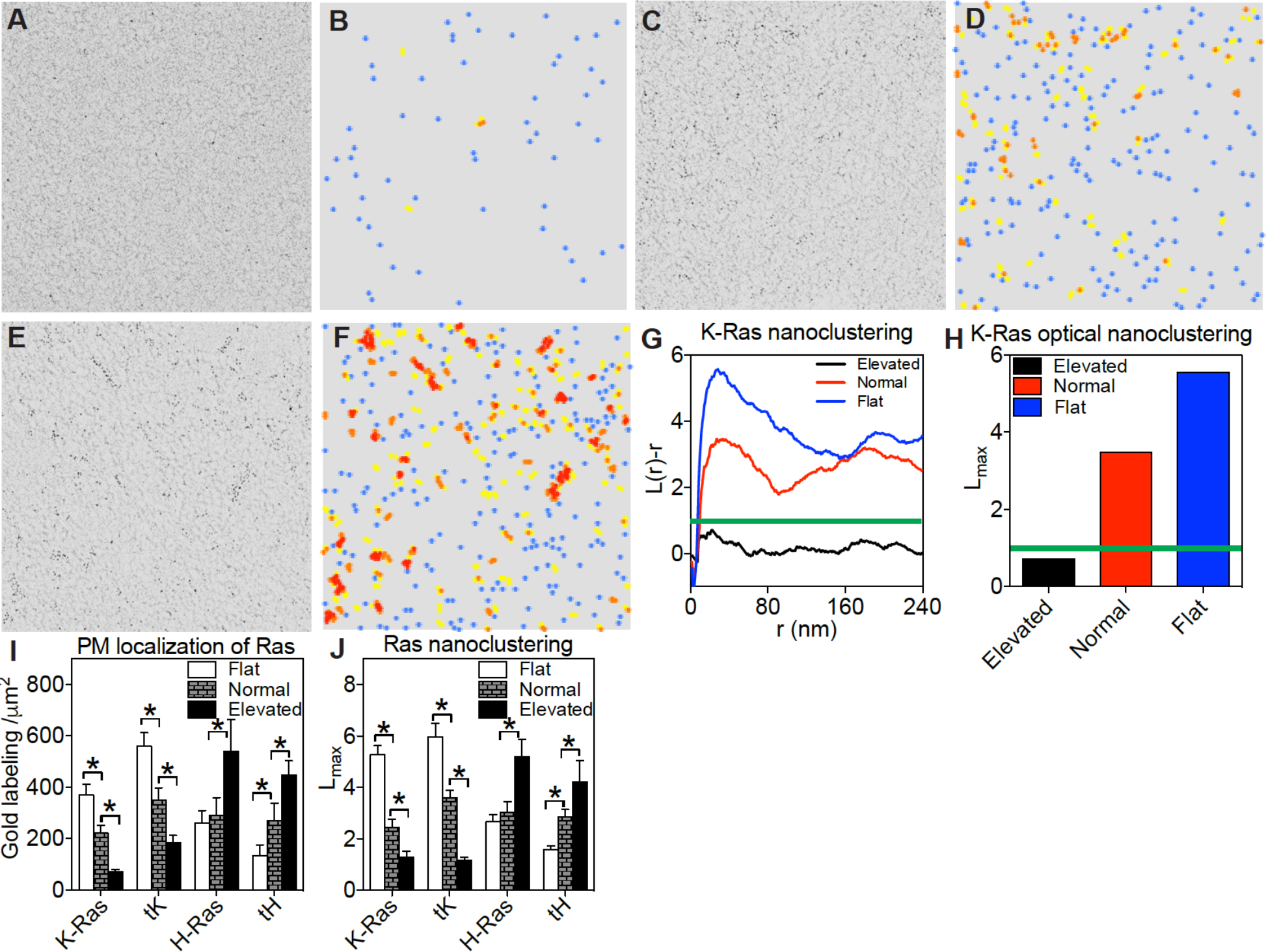
Electron microscopy (EM) univariate spatial analysis characterizes the spatiotemporal organization of Ras isoforms on the PM. Baby hamster kidney (BHK) cells expressing the GFP-K-Ras^*G12V*^ were subjected to various manipulations to modulate local PM curvature before their intact apical PM sheets were attached to EM grids and immunogold labeling. An electron micrograph of an intact PM sheet of BHK cells elevated PM curvature (**A**) is shown, along the corresponding heatmap showing the color-coded spatial distribution of the same gold nanoparticles (**B**). An EM image of gold-labeled GFP-K-Ras^*G12V*^ in a BHK cell with normal PM curvature (**C**) and its corresponding distribution heatmap (**D**) are shown. Additionally, an EM image of gold-labeled GFP-K-Ras^*G12V*^ in a BHK cell with elevated local PM curvature (**E**) and its corresponding distribution heatmap (**F**) are also shown. The color-coding indicates the spatial distribution: blue dots mean monomers, yellow dots mean dimers, orange dots indicate trimers and red dots indicate higher ordered multimers. (**G**) The extent of nanoclustering, *L*(*r*)-*r*, was plotted against the cluster radius, *r*, in nanometers. The *L*(*r*)-*r* values above the 99% confidence interval (99%C.I.) indicate statistically meaningful clustering, with the larger *L*(*r*)-*r* values indicating more extensive clustering. (**H**) The peak values of *L*(*r*)-*r* function curves, termed as *L*_*max*_, were used to summarize the clustering data. The PM localization (**I**) and the nanoclustering (**J**) of Ras proteins/peptides are shown as a function of gradual increase in PM curvature. Data in (I) and (J) are shown as mean±SEM. The statistical significance in gold numbers was evaluated using one-way ANOVA with * indicating p < 0.05. The statistical significance of the nanoclustering was evaluated using the bootstrap tests via ranking against 1000 Monte Carlo simulations, with * indicating p < 0.05.

**Supplemental Figure S2.**
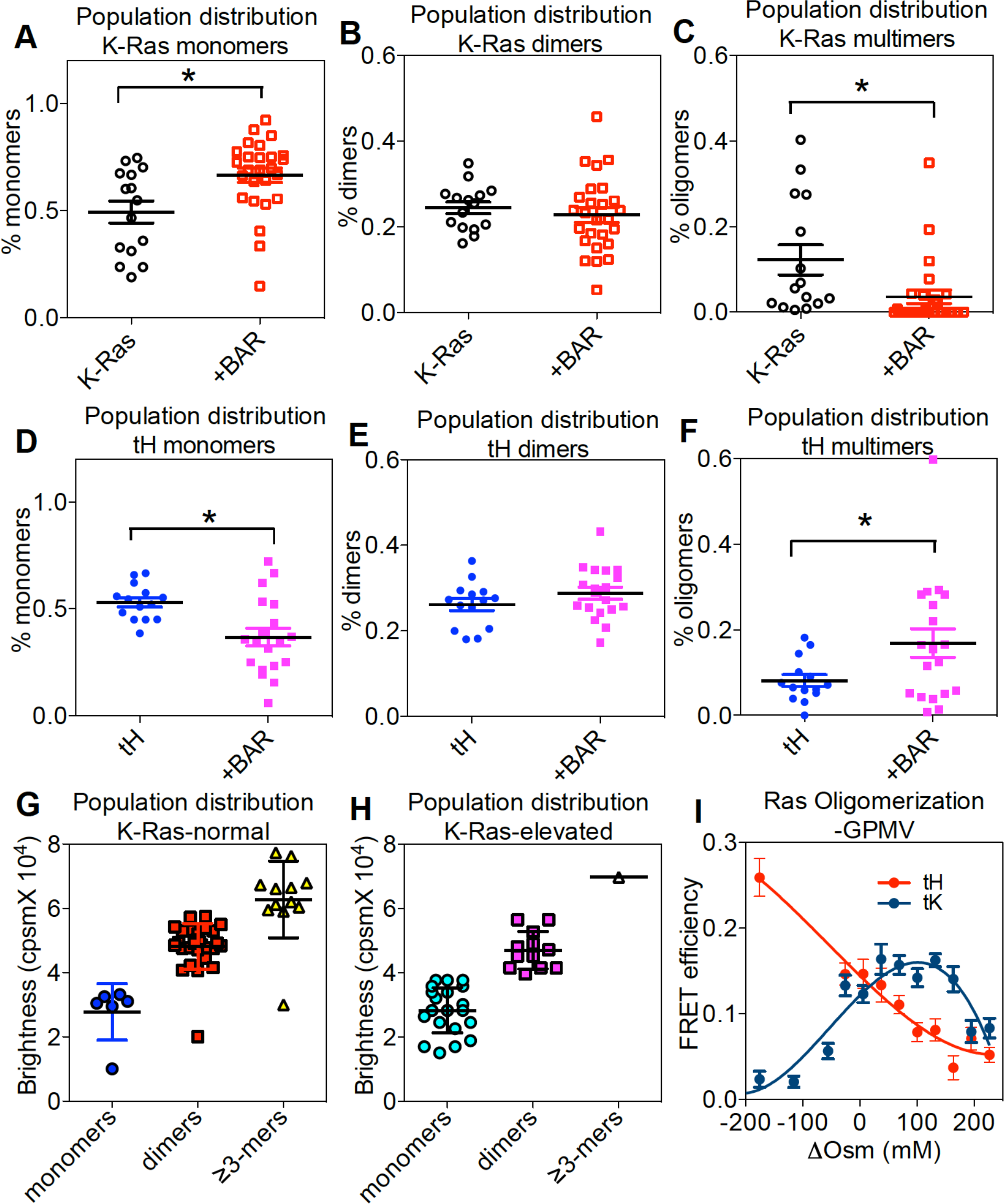
Supplemental Figure S2. PM curvature-induced changes in the oligomerization of Ras isoforms on the PM. The EM spatial data were further interrogated. Population distribution of GFP-K-RasG12V monomers (**A**), dimers (**B**) and oligomers (**C**), or GFP-tH monomers (**D**), dimers (**E**) and oligomers (**F**) was counted. In a RICS-B/N analysis, the spatial aggregation of GFP-K-RasG12V was studied in BHK cells with various local PM curvature. Specifically, the fluorescence intensities of GFP-K-RasG12V were measured on the PM of live BHK cells and used to quantify the oligomerization of GFP-K-RasG12V. GFP fluorescence brightness was plotted a number of molecules within an aggregate in BHK cells with normal PM curvature (untreated cells, **G**) or with elevated PM curvature (ectopic expression of RFP-BAR_amph2_ domain, **H**). (**I**) The extent of Ras oligomerization as a function of local membrane curvature fluctuation was evaluated using FLIM-FRET in GPMVs. GFP fluorescence lifetime was measured and was used to calculate FRET efficiency as a function of ΔOsm. For panels A-H, all individual data points are shown. For panel I, data are shown as mean±SEM. The statistical significance was evaluated using one-way ANOVA with * indicating p < 0.05.

**Supplemental Figure S3.**
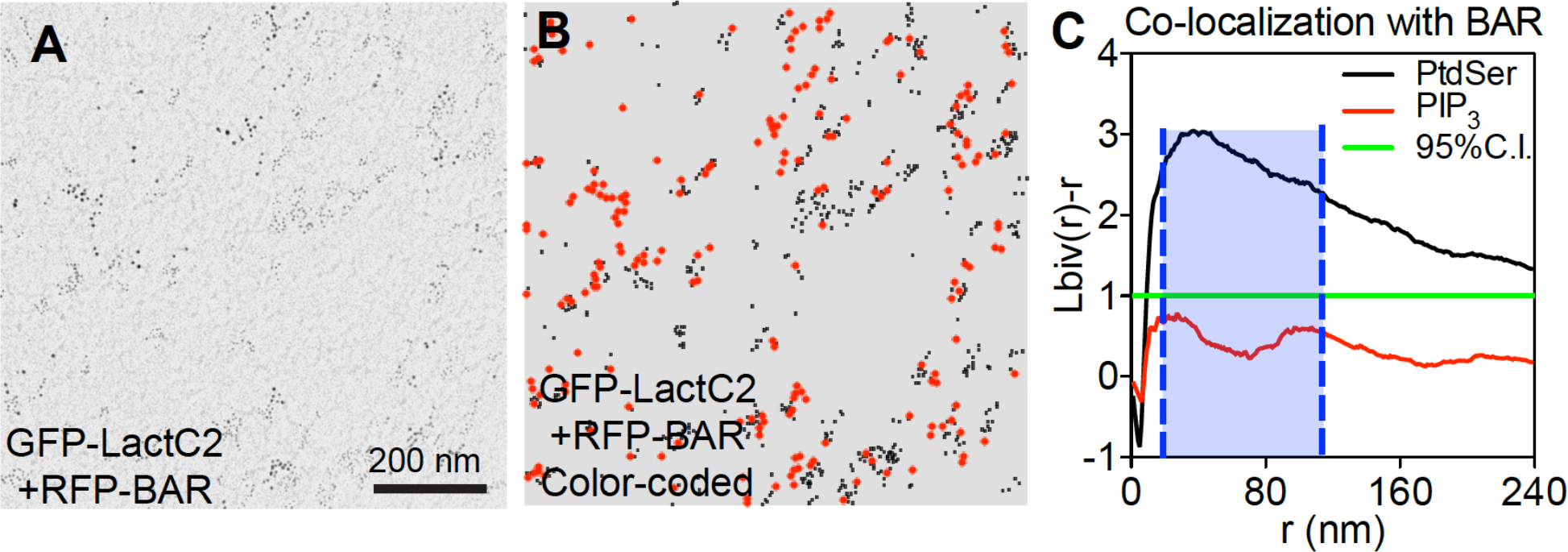
Bivariate co-clustering analysis quantifies PM curvature preferences of various lipids in the PM. (***A***) Image of an intact PM sheet of 1µm^2^ area of BHK cells co-expressing GFP-LactC2 and RFP-BAR_amph2_ domain under the isotonic condition is shown. The PM sheet was immunogold co-labeled with the 6nm gold conjugated to anti-GFP antibody and the 2nm gold conjugated to anti-RFP antibody, respectively. Gold distribution was imaged using TEM at 100,000X magnification. (***B***) Gold nanoparticles in the EM image in (A) are color-coated to clarify the two populations of gold: red and big dots = 6nm gold labeling GFP; black and small dots = 2nm gold labeling RFP. (***C***) The extent of co-clustering, *L*_*biv*_(*r*)-*r*, was plotted against the cluster radius, *r*, in nanometers. The *L*_*biv*_(*r*)-*r* values above the 95% confidence interval (95%C.I.) indicate statistically meaningful co-localization between the two populations of gold, with the larger *L*_*biv*_(*r*)-*r* values indicating more extensive co-localization. Each *L*_*biv*_(*r*)-*r* curve was integrated between 10 and 110nm distance (blue shade) and yield an LBI value to summarize the extent of co-localization. The statistical significance was evaluated using the bootstrap tests via ranking against 1000 Monte Carlo simulations.

**Supplemental Figure S4.**
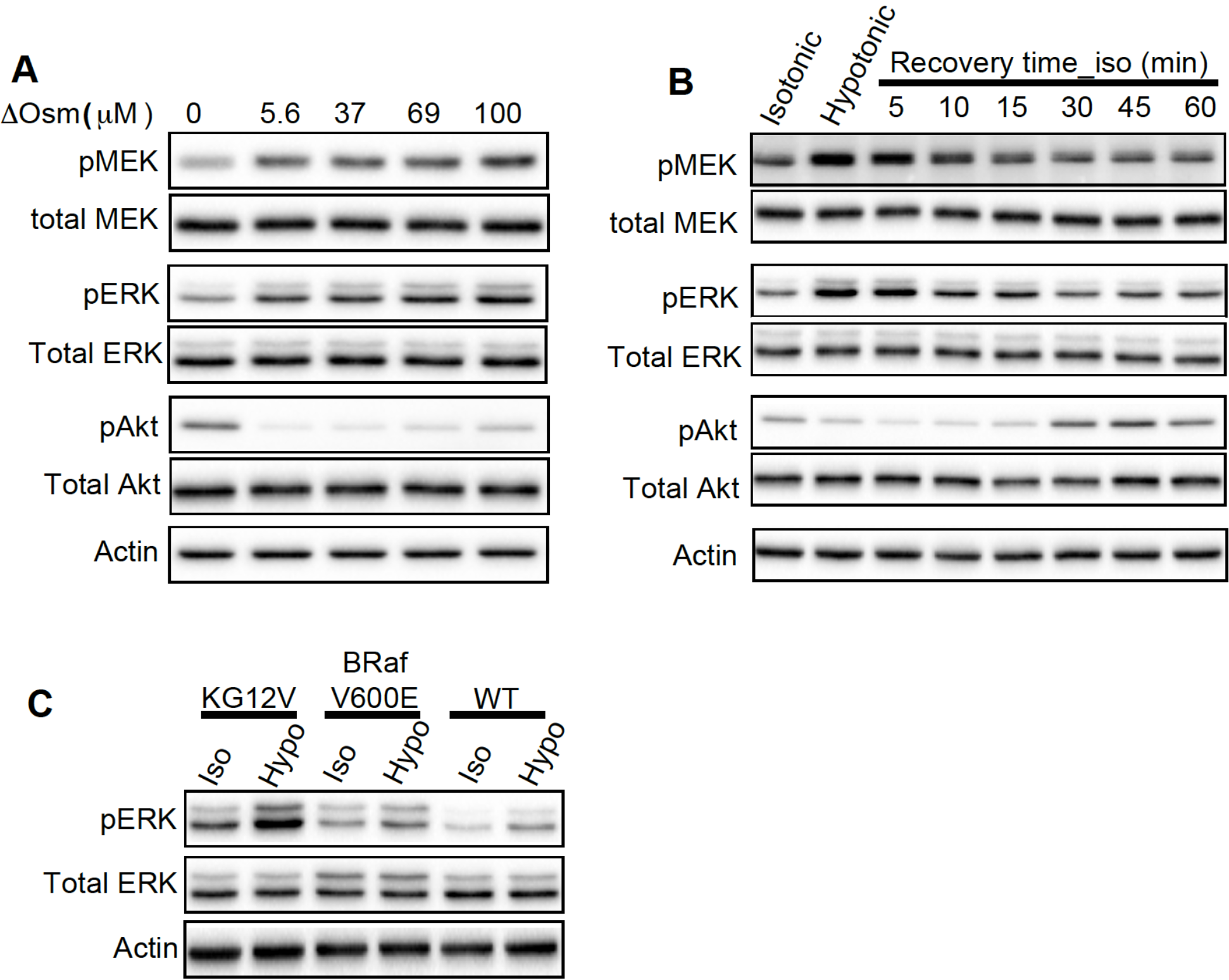
Ras-dependent MAPK and PI3K signaling is sensitive to PM curvature modulations. (**A**) Sample immunoblots of MAPK and PI3K signaling in pre-serum-starved BHK cells subjected to the media with various osmolarity for 5 minutes are shown. (**B**) Pre-serum-starved BHK cells were incubated in the media containing 40% water for 5 minutes before being returned to the isotonic media for various time periods. (**C**) Ras-less MEF lines were subjected to HEPES buffers at either isotonic (Iso) or hypotonic (Hypo) osmolalities for 5 minutes before harvesting. The whole cell lysates were used for blotting against the phosphorylated MEK, ERK and Akt (pMEK, pERK and pAkt). Three independent experiments were conducted for each set.

